# TOR coordinates nucleotide availability with ribosome biogenesis in plants

**DOI:** 10.1101/2020.01.30.927418

**Authors:** Michael Busche, M. Regina Scarpin, Robert Hnasko, Jacob O. Brunkard

## Abstract

TARGET OF RAPAMYCIN (TOR) is a conserved eukaryotic Ser/Thr protein kinase that coordinates growth and metabolism with nutrient availability. We conducted a medium-throughput functional genetic screen to discover essential genes that promote TOR activity in plants, and identified a critical regulatory enzyme, cytosolic phosphoribosyl pyrophosphate (PRPP) synthetase (PRS4). PRS4 synthesizes cytosolic PRPP, a key upstream metabolite in nucleotide synthesis and salvage pathways. We found that *prs4* knockouts are embryo-lethal in *A. thaliana*, and that silencing *PRS4* expression in *N. benthamiana* causes pleiotropic developmental phenotypes, including dwarfism, aberrant leaf shape, and delayed flowering. Transcriptomic analysis revealed that ribosome biogenesis is among the most strongly repressed processes in *prs4* knockdowns. Building on these results, we discovered that TOR activity is inhibited by chemical or genetic disruption of nucleotide biosynthesis, but that this effect can be reversed by supplying plants with physiological levels of nucleotides. Finally, we show that TOR transcriptionally promotes nucleotide biosynthesis to support the demands of ribosomal RNA synthesis. We propose that TOR coordinates ribosome biogenesis with nucleotide availability in plants to maintain metabolic homeostasis and support growth.

## Introduction

Phosphorus (P) is an essential element for plant life, but is not highly available in agricultural soils, and is the primary limiting factor for crop yield on more than 30% of global arable land (Vance et al., 2003; López-Arredondo et al., 2014). Crop fertilizers include high levels of phosphate (P_i_) to drive plant growth and increase crop yields, but crops only utilize ∼20-30% of P_i_ applied to soils, a significant waste of resources that is also environmentally harmful (López-Arredondo et al., 2014; Correll, 1998). Moreover, P_i_ mineral deposits are finite, with P_i_ production from mining predicted to begin declining within fifteen years (Cordell and White, 2011), followed by global depletion of P_i_ mineral deposits within as little as one century. Current models suggest that P_i_ fertilizer use will exceed safe planetary boundaries by 2050 without significant technological advances to reduce the need for P_i_ fertilizers (Springmann et al., 2018). Therefore, a major agronomic goal is to maximize P_i_ uptake and utilization, including by breeding and gene editing to rewire P_i_ sensing, signaling, and allocation networks in crop species.

Approximately 50% of the organic phosphorus (P_o_) in leaves is incorporated in ribosomal RNA (rRNA) (Veneklaas et al., 2012), with the remaining P_o_ divided among other RNAs, DNA, phospholipids, and P-esters (phosphorylated metabolites, such as glucose-6-phosphate and free nucleotides, as well as phosphorylated proteins) (Figure 1). Since the majority of P_o_ is invested in ribosomes, some species in the *Proteaceae* family have adapted to P_i_-poor soils by synthesizing relatively few ribosomes, with as little as 40-fold less ribosomes per leaf fresh weight when compared to *Arabidopsis thaliana* (Sulpice et al., 2014). While reducing the number of ribosomes prevents P_i_ deprivation stress in these plants, the consequential decrease in the rate of protein synthesis dramatically slows plant growth (Sulpice et al., 2014). Balancing the trade-off between P_i_ demand and growth rates by genetically optimizing ribosome abundance in different tissues and developmental stages in crop species is a promising strategy to reduce agricultural reliance on P_i_ fertilizers (Veneklaas et al., 2012). Achieving this goal will require deep understanding of the regulation of ribosome biogenesis and the P_i_- and P_o_-sensing mechanisms in plants.

**Figure 1.**
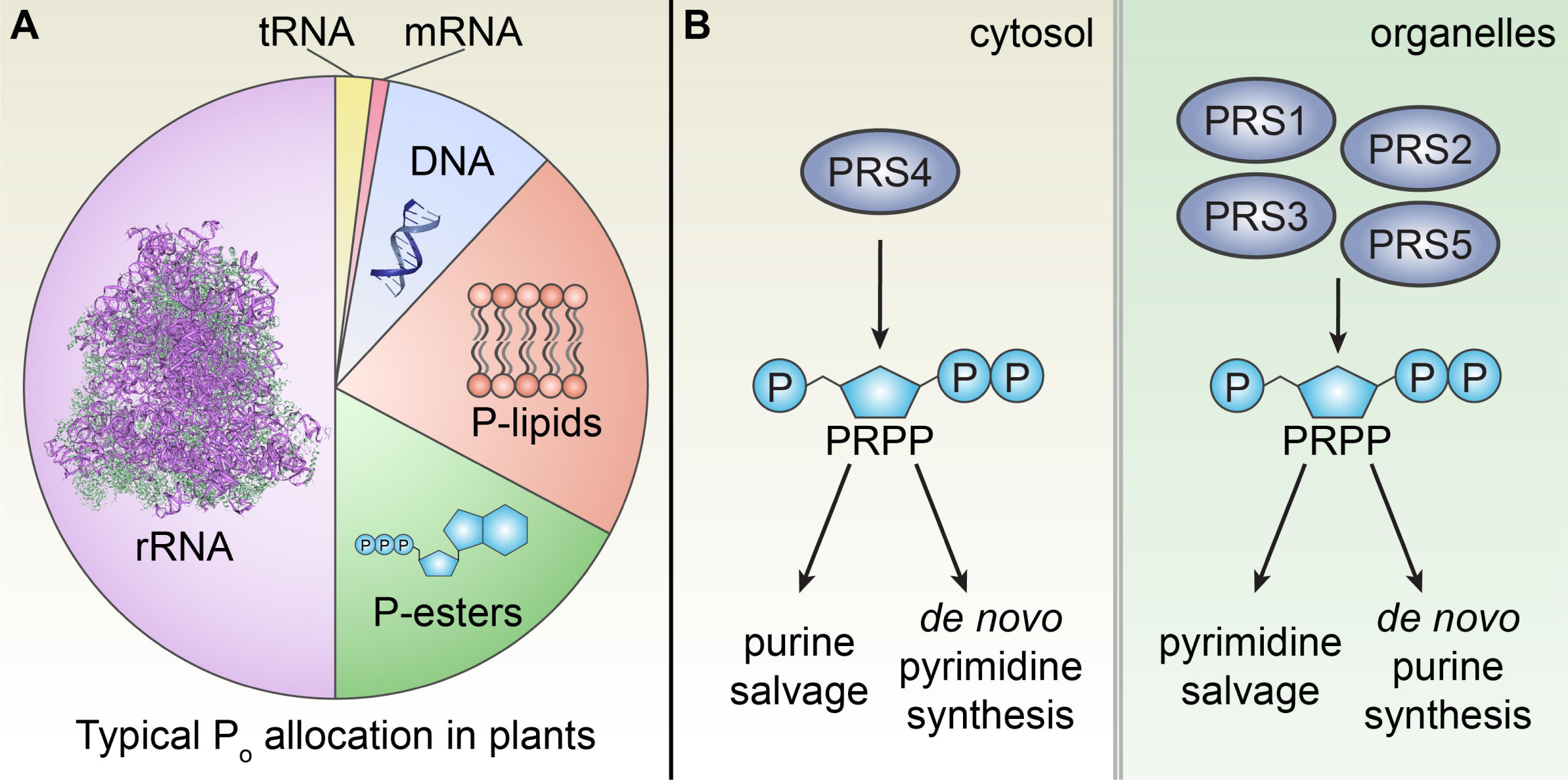
Ribosomes are major sinks for organic phosphorus in plant cells. **(A)** As much as 50% of organic phosphorus (P_O_) is allocated to ribosomal RNA in plant cells. The remaining P_o_ is allocated to nucleic acids (transfer RNAs (∼2%), messenger RNAs (∼1%), and DNA (∼7%)), phospholipids (∼30%), and phosphoesters (∼20%), including phosphorylated proteins and metabolites, e.g., free nucleotide triphosphates. **(B)** Plant genomes encode several phosphoribosyl pyrophosphate (PRPP) synthetases (PRSs) that localize to either the cytosol (left) or organelles (right). In *A. thaliana*, there are five *PRS* genes: *PRS4* encodes the only cytosolic PRS, and *PRS1, PRS2, PRS3*, and *PRS5* encode organelle-targeted PRSs. In the cytosol, PRPP is primarily metabolized via the purine salvage and *de novo* pyrimidine synthesis pathways. Conversely, in the organelles, PRPP is metabolized in the pyrimidine salvage and *de novo* purine synthesis pathways.

TARGET OF RAPAMYCIN (TOR) is a conserved eukaryotic kinase that broadly coordinates metabolism with nutrient availability (Chantranupong et al., 2015; Shi et al., 2018), and is a crucial regulator of ribosome abundance (Hannan et al., 2003; Delarue et al., 2018). When nutrients are available, TOR is active and promotes anabolism and growth; when nutrients are scarce, TOR is inactive and cells become quiescent, using catabolic pathways for sustenance (Saxton and Sabatini, 2017). Mutations in the TOR signaling network, especially mutations that increase TOR activity, can cause or contribute to a wide range of cancers (Grabiner et al., 2014; Saxton and Sabatini, 2017). One consequence of elevated TOR activity in mutated human cell lines is increased ribosome biogenesis, because TOR strongly promotes the expression of ribosomal protein genes, rRNA, and other genes that contribute to ribosome assembly (Pelletier et al., 2017). As a result, cells with high TOR activity have high nucleotide demands, and are hypersensitive to chemotherapeutic nucleotide biosynthesis inhibitors (Valvezan et al., 2017; Vander Heiden and DeBerardinis, 2017). To meet these nucleotide demands, TOR promotes pyrimidine and purine biosynthesis (Ben-Sahra et al., 2013; Robitaille et al., 2013; Ben-Sahra et al., 2016), effectively reorganizing cellular metabolism to support higher levels of rRNA synthesis. Recent studies have further demonstrated that TOR activity is coordinated with nucleotide availability in human cells (Emmanuel et al., 2017; Hoxhaj et al., 2017). This coordination is proposed to be mediated by GTP levels: under purine-limiting conditions, GTP levels decrease, disabling the small GTPase Rheb that promote TOR activity in humans (Emmanuel et al., 2017; Hoxhaj et al., 2017). Thus, TOR senses nucleotide availability to serve as a metabolic checkpoint that maintains homeostasis by coupling nucleotide demand (primarily rRNA synthesis) with nucleotide biosynthesis.

Nucleotide metabolism and signaling are well-studied in humans, because nucleotide homeostasis is often dysregulated in human diseases (Aird and Zhang, 2015), but little is known about how plants sense and respond to nucleotides. P_i_ deprivation can cause a five-fold reduction in free nucleotide levels in plants, making nucleotides a potentially limiting metabolite in soils with poor P_i_ availability (Theodorou and Plaxton, 1993). In *Chlamydomonas reinhardtii*, a photosynthetic algal species, P_i_ starvation reduces TOR activity and genetic manipulation of P_i_ metabolism disrupts TOR signaling, demonstrating that TOR can contribute to the maintenance of P_i_ homeostasis (Cuoso *et al*., 2020). A key intermediate in the incorporation of P_i_ into nucleotides is phosphoribosyl pyrophosphate (PRPP). PRPP is synthesized by phosphoribosyl pyrophosphate synthetase (PRS, sometimes called ribose-phosphate diphosphokinase), which transfers the diphosphoryl group from ATP to ribose-5-phosphate (the end product of the pentose phosphate pathway) in a Mg^2+^-dependent reaction. PRPP then acts as a donor of phospho-ribose groups for nucleotide and amino acid synthesis, among other biochemical pathways. PRS has emerged as a key regulatory enzyme in human cellular nutrient signaling pathways; for example, PRS-mediated synthesis of PRPP is the rate-limiting step that drives the protein and nucleotide synthesis necessary for tumorigenesis downstream of the Myc oncogene (Cunningham et al., 2014).

Plant genomes encode several PRS enzymes, with at least one localized to the plastid and another localized to the cytosol (Figure 1B). Plastids have spatially co-opted many of the PRPP-dependent metabolic pathways in plants, including *de novo* purine, histidine, and tryptophan biosynthesis, the pyrimidine salvage pathway, as well as various metabolic pathways downstream of these molecules, including cytokinin biosynthesis. Cytosolic PRPP is therefore primarily required to regenerate nucleotides via the purine salvage pathway and for *de novo* pyrimidine biosynthesis (Lerbs et al., 1980; Witz et al., 2012). In *Arabidopsis thaliana*, four genes encode organellar PRS enzymes (*PRS1, PRS2, PRS3,* and *PRS5*) and one gene encodes the cytosolic PRS enzyme (*PRS4*) (Ito et al., 2011). Here, we show that *PRS4* is an essential gene in plants that is required to maintain TOR activity. Furthermore, taking a functional genetics approach in *N. benthamiana* and a chemical genetic approach in *A. thaliana*, we specifically demonstrate that nucleotides promote TOR activity, and that TOR transcriptionally promotes nucleotide biosynthesis. We conclude that TOR coordinates nucleotide availability with ribosome biogenesis to maintain P homeostasis in plants.

## Results

### *PRS4* drives TOR activity and plant development

In an effort to identify genes that regulate TOR activity, we conducted a medium-throughput functional genetic screen using virus-induced gene silencing (VIGS) to silence *N. benthamiana* genes that we predicted to be essential for embryogenesis in *A. thaliana*. To measure TOR activity after gene silencing, we extracted leaf proteins to prepare Western blots and probed these blots with α-AtS6K-p449, which specifically detects a phosphopeptide on plant S6 KINASE (S6K) that is directly phosphorylated by the TOR kinase complex (Xiong and Sheen, 2012). We found that silencing *PRS4* strongly reduces S6K-pT449 levels compared to mock treatment (Figure 2A). We then probed total S6K protein levels with α-S6K, confirming that total S6K levels do not decrease in TRV::*NbPRS4*, and therefore demonstrating that TOR activity is severely reduced in TRV::*NbPRS4* knockdown leaves (Figure 2A).

**Figure 2.**
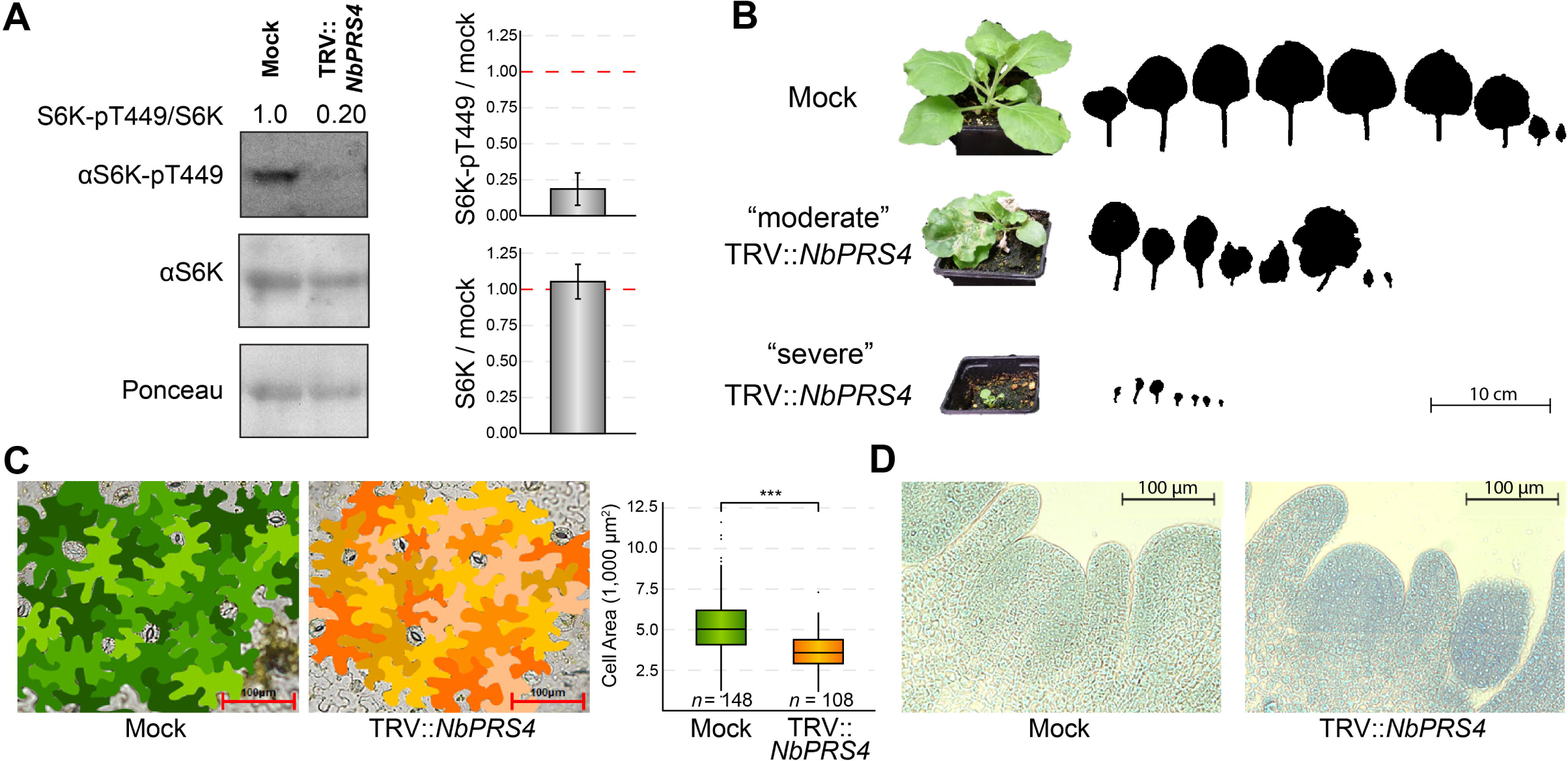
Cytosolic PRS (PRS4) drives plant development and TOR activity in *N. benthamiana*. **(A)** Silencing *PRS4* drastically reduces TOR activity. S6K-pT449 levels reflect TOR activity, because S6K-T449 is a direct substrate of TOR (Xiong and Sheen, 2012). S6K-pT449 and total S6K levels were assayed by Western blots in knockdown TRV::*NbPRS4* plants or mock controls (representative images shown here). Quantification of band densities confirmed that S6K-pT449 levels decrease ∼5-fold in TRV::*NbPRS4* knockdowns, but total S6K levels are not affected in TRV::*NbPRS4*. **(B)** *PRS4* is required for shoot development. There are fewer leaves in TRV::*NbPRS4* knockdowns, and the leaves are misshapen and small. We observed individual-to-individual variation in phenotypic severity after silencing *PRS4* by VIGS; a representative of the “moderate” TRV::*NbPRS4* phenotype and of the “severe” TRV::*NbPRS4* phenotype are shown. Outlines of leaf shapes are shown, including all leaves with silenced *PRS4* expression (i.e., only leaves above the primary infected leaf), with the oldest leaf on the left and the youngest leaf on the right. **(C)** Silencing *PRS4* impairs cell expansion and cell division. Epidermal pavement cell shape was not dramatically altered in TRV::*NbPRS4* knockdowns, but epidermal pavement cell size was significantly lower. This difference in cell size is insufficient to account for the decrease in total leaf area shown in panel B; therefore, there are also fewer epidermal pavement cells in TRV::*NbPRS4*. **(D)** We did not observe clear effects of silencing *PRS4* on vegetative shoot apical meristem morphology.

Knocking down *PRS4* expression with TRV::*NbPRS4* caused a range of dramatic developmental phenotypes (Figure 2B), including delayed flowering, misshapen leaves, dwarfism, and even lethality: 14% of TRV::*NbPRS4* knockdown plants died shortly after *PRS4* silencing was induced (*n* = 50), whereas none of the mock-treated plants died. Surviving TRV::*NbPRS4* leaves were misshapen and as small as 5% the size of leaves in mock-treated shoots (Figure 2B). Epidermal cells were significantly smaller in TRV::*NbPRS4* plants (71% the size of cells in mock-treated leaves, *n* ≥ 108, *p* < 10^-3^, Figure 2C), but the difference in cell size was not enough to explain the severe reduction in leaf size (Figure 2B), suggesting that *PRS4* is required to promote cell expansion and cell division. In TRV::*NbPRS4* plants, developmental timing of leaf initiation and flowering was severely disrupted. Leaf initiation was delayed or completely arrested three weeks after infiltrating VIGS vectors in TRV::*NbPRS4* plants (Figure 2B). TRV::*NbPRS4* plants flowered two or more weeks later than mock-treated plants, and many TRV::*NbPRS4* knockdowns did not flower at all during our six-week observation period. Despite these clear defects in shoot development, we did not detect any striking morphological phenotype in histological sections of TRV::*NbPRS4* knockdown vegetative shoot apical meristems (Figure 2D). Taken together, these results show that *PRS4* is required for diverse developmental processes, including leaf initiation, expansion, and morphology; the transition from vegetative to reproductive growth; and both cell expansion and cell division.

### *PRS4* is essential for plant embryogenesis

As part of a recent high-throughput effort to generate higher order mutations in *A. thaliana*, *prs4*/+ was crossed to *prs3/*+, but no offspring in the F_2_ generation of this cross were homozygous for *prs4*, hinting that *PRS4* is essential for *A. thaliana* development (Bolle et al., 2013). To test this hypothesis, we obtained a putative T-DNA insertion line in the *PRS4* gene (*At2g42910*), GabiKat-780B11 (Figure 3). Preliminary sequencing suggested that GabiKat-780B11 carries a T-DNA insertion in the third intron of the *PRS4* gene, and is therefore predicted to be a null allele. No other publicly available T-DNA insertion lines are predicted to abolish *PRS4* expression. We confirmed the location of the GabiKat-780B11 T-DNA by PCR (Figure 3A), and named this allele *prs4-1.* We isolated several independent *prs4-1*/+ heterozygous plants, allowed the plants to self-fertilize, collected seed, and genotyped the F_1_ generation. We identified only *prs4-1/*+ heterozygous or +/+ plants in a 2:1 ratio, indicating that *prs4-1/prs4-1* is lethal.

**Figure 3.**
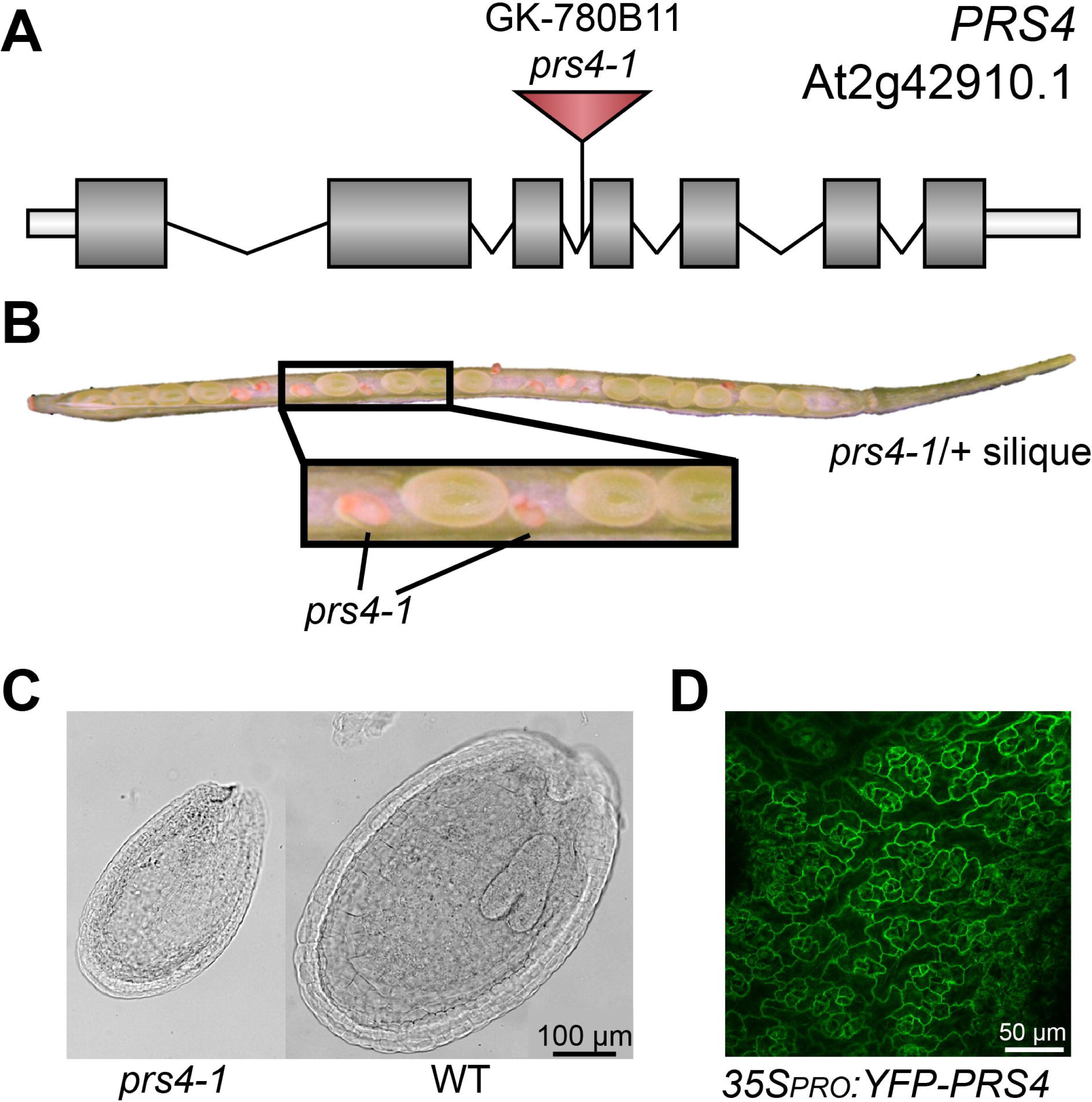
Cytosolic PRS4 is required for embryogenesis. **(A)** *PRS4* is encoded by *At2g42910*. We isolated an insertional allele, GK-780B11, that we named *prs4-1*. **(B)** Representative silique of a self-fertilized *prs4-1* / + heterozygous parent. 23% of seeds are *prs4-1 / prs4-1* homozygotes; these seeds are shrunken and brown. **(C)** Homozygous *prs4-1* seeds have no readily visible embryo after clearing. A sibling WT seed from the same silique with a clearly visible early torpedo-shaped embryo is shown for comparison. **(D)** *35S_PRO_:YFP-PRS4* expressed in *A. thaliana* localizes to the cytoplasm.

To confirm that *prs4-1* is lethal and determine when *prs4-1* mutants arrest, we next examined seeds in the siliques of several *prs4-1*/+ plants from separate families. 77% of seeds in these siliques were green with viable embryos, whereas 23% of these seeds were collapsed and brown (Figure 3B). After histochemically clearing the seeds, we observed normal embryo morphology in the green seeds, but could not observe a multicellular embryo in the shrunken seeds (Figure 3C). The green seeds segregated 2:1 for *prs4-1/+* : +/+, as predicted for an embryo-lethal allele. We marked the position of putative *prs4-1/prs4-1* aborted seeds, and noticed that the expected number of seeds near the apex of the silique were aborted (26.7%, *n* = 180, *χ^2^* = 0.3, *p* = 0.61), but less than 20% of seeds near the base of the silique were aborted (18.9%, *n* = 180, *χ^2^* = 3.6, *p* = 0.058), suggesting a slight pollen transmission defect that could explain why only 23% of *prs4-1*/+ offspring are homozygous for the null *prs4-1* allele (Meinke, 1985). Finally, we generated stable transgenic lines by transforming *prs4-1*/+ flowers with T-DNAs carrying either *35S_PRO_:PRS4-GFP* or *35S_PRO_:YFP-PRS4* (Figure 3D), both of which rescued the lethal phenotype of *prs4-1/prs4-1* seeds. Therefore, we concluded that *PRS4* is required for the earliest stages of *A. thaliana* sporophyte development.

In wild-type *A. thaliana*, *PRS4* transcriptional expression is strongest in metabolically active cells, especially the shoot apical meristem and germinating seeds (Supplemental Figure 1). *PRS4* transcriptional expression is notably lower in quiescent plants, such as seeds entering dormancy and mature pollen grains (Supplemental Figure 1). The expression patterns of genes that encode the organelle-targeted PRS enzymes, *PRS2*, *PRS3*, and *PRS5*, are largely similar to *PRS4*, although *PRS2* is expressed in seeds entering dormancy, and both *PRS2* and *PRS3* are expressed in mature pollen grains (Supplemental Figure 1). *PRS1* is consistently expressed at very low levels during development, in contrast to the other *PRS* genes (Supplemental Figure 1). These expression patterns indicate that PRS enzymes are strongly expressed in both the cytosol and the organelles in all metabolically active developmental stages to support PRPP synthesis.

### *PRS4* promotes ribosome biogenesis

To identify disrupted genetic pathways in TRV::*NbPRS4* plants, we took a global transcriptomic approach (Figure 4). Nine RNA-Seq libraries were sequenced from rRNA-depleted RNA extracted from replicated pools of *N. benthamiana* shoot apices (defined as all tissue above the second-youngest leaf, including the shoot meristem, some stem, and the youngest two visible leaves; Supplemental Figure 2). Three libraries each were prepared from mock-silenced plants, TRV::*NbPRS4* plants with “moderate” visible phenotypes (Figure 2B, middle panel), or TRV::*NbPRS4* plants with “severe” visible phenotypes (Figure 2B, bottom panel). RNA-Seq confirmed that *PRS4* expression is effectively silenced in the TRV::*NbPRS4* plants, but not in mock-infected plants, as expected. 35,877 annotated *N. benthamiana* genes were detected in our transcriptomes, of which 4,986 were significantly differentially-expressed genes (DEGs) between TRV::*NbPRS4* and mock-silenced plants (3,213 induced, 1,773 repressed, *p_adj_.* < 0.05) (Figure 4A, Supplemental Data Sets 1 and 2). This large number of DEGs demonstrates the dramatic effects of disrupting PRS4 activity on cellular homeostasis. We detected relatively few DEGs between TRV::*NbPRS4* plants with “moderate” or “severe” visible phenotypes (489 genes, *p_adj_* < 0.05) (Figure 4A, Supplemental Data Set 3), suggesting that the morphological variation among TRV::*NbPRS4* knockdown plants does not strongly correlate with differences in gene expression. Principal components analysis of the nine transcriptomes also showed that the mock-silenced transcriptomes grouped separately from the TRV::*NbPRS4* knockdown transcriptomes, but that the “moderate” and “severe” groups were not so readily distinguished (Figure 4A).

**Figure 4.**
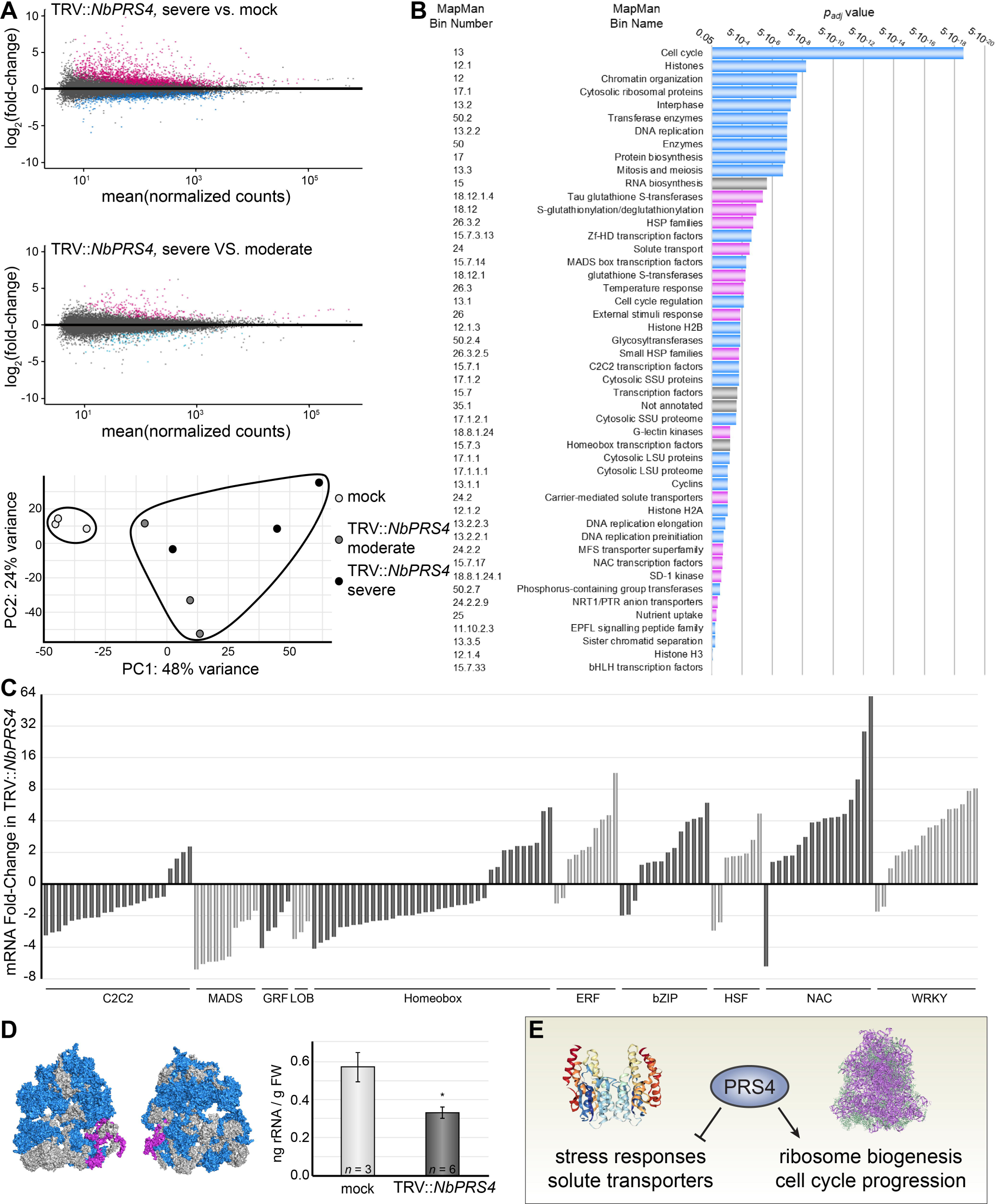
Silencing PRS4 reprograms the transcriptome to repress ribosome biogenesis. **(A)** Scatterplots of gene expression changes in *N. benthamiana* after VIGS. 4,986 genes were significantly differentially expressed between TRV::*NbPRS4* knockdowns with severe phenotypes and mock plants (top panel), but only 489 genes were significantly differentially expressed between TRV::*NbPRS4* knockdowns with severe versus moderate phenotypes (middle panel). Principal component analysis demonstrates that the mock-treated transcriptomes are readily distinguished from the TRV::*NbPRS4* knockdowns, but that the TRV::*NbPRS4* transcriptomes from plants with severe or moderate phenotypes are not grouped separately. **(B)** MapMan functional analysis of DEGs in TRV::*NbPRS4* revealed 48 significantly-affected categories (*p_adj_*. < 0.05). Categories with primarily repressed gene expression are indicated in blue; categories with primarily induced gene expression are indicated in magenta; categories with genes that are both induced and repressed (e.g., transcription factors) are shown in grey. **(C)** Transcription factor mRNA levels are significantly different in TRV::*NbPRS4* knockdowns. Select transcription factor families typically involved in plant development (C2C2, Cys_2_Cys_2_ Zn-finger family; MADS, MCM1/AGAMOUS/DEFICIENS/SRF box family; GRF, GROWTH-REGULATING FACTOR family; LOB, LATERAL ORGAN BOUNDARIES family; and the homeobox family) or plant stress responses (ERF, ETHYLENE RESPONSE FACTOR family, excluding AP2 orthologues; bZIP, basic Leucine Zipper Domain family; HSF, Heat Shock Factor; NAC, NAM/ATAF1/CUC2 family; and the WRKY family). **(D)** mRNAs encoding 51 subunits of the plant cytosolic ribosome were down-regulated in TRV::*NbPRS4* (shown in this model of a ribosome in blue); mRNAs encoding 2 subunits were up-regulated (magenta). There was ∼50% less total rRNA in TRV::*NbPRS4* knockdowns (\**p* < 0.05). **(E)** Transcriptomic analysis demonstrates that PRS4 is required to promote expression of genes that contribute to ribosome biogenesis (purple, right) and cell cycle progression, and to repress induction of starvation and oxidative stress response genes, including solute transporters and antioxidant glutathione S-transferases (rainbow, left).

*N. benthamiana* is an emerging model system whose genome has not been extensively annotated, so to develop tools to rigorously analyze the TRV::*NbPRS4* RNA-Seq data, we submitted the entire *N. benthamiana* predicted proteome (genome version 1.0.1) to Mercator4 for functional annotation using the MapMan gene ontology (Lohse et al., 2014). This approach conservatively assigned functions to 23,001 annotated genes in *N. benthamiana* (Supplemental Data Set 3), of which 14,502 were detected in our RNA-Seq analysis and 2,662 were DEGs (Supplemental Data Set 1). We then used MapMan to identify enriched functional categories in our transcriptomes, with stringent parameters, using only significant DEGs and the Benjamini-Hochberg-Yekutieli procedure to correct for the false discovery rate. 48 categories were significantly affected in TRV::*NbPRS4* versus mock (*p_adj_* < 0.05) (Figure 4B, Supplemental Data Set 5), but no categories were significantly affected in the comparison between the visually “severe” and “mild” phenotype pools of TRV::*NbPRS4* plants (*p_adj_* < 0.05), again supporting the hypothesis that the morphological variation we observed was not due to a consistent difference in the plants’ transcriptional programs.

In the TRV::*NbPRS4* transcriptome, one of the most significant effects was downregulation of genes involved in ribosome biogenesis (Figure 4D, Supplemental Data Sets 1, 5). To illustrate, the expression of genes encoding 51 of the 81 plant cytosolic ribosomal proteins were significantly lower in TRV::*NbPRS4* than in mock controls (Figure 4D, Supplemental Data Set 4). TOR has conserved roles in promoting expression of ribosomal protein genes (Cardenas et al., 1999; Hannan et al., 2003; Marion et al., 2004; Xiong et al., 2013a), so their repression likely reflects the suppression of TOR activity in TRV::*NbPRS4* plants. To determine the consequences of ribosome biogenesis transcript downregulation in TRV::*NbPRS4* on overall ribosome abundance, we assayed total rRNA in TRV::*NbPRS4* versus mock plants, and found that rRNA levels were approximately 2-fold lower per fresh weight in TRV::*NbPRS4* knockdown shoots than in mock-treated plants (*n* ≥ 3, *p* < 0.05, Figure 4D). Since ribosomes can account for ∼60% of nucleotides and ∼25% of proteins in a plant cell, a two-fold reduction in ribosome abundance represents a dramatic shift in cellular physiology. Thus, a major effect of knocking down *PRS4* expression is disruption of ribosome biogenesis, likely a consequence of TOR inactivation (Figure 4E).

Various other biological processes were also significantly affected in the TRV::*NbPRS4* transcriptomes (Figure 4B). Genes involved in cell cycle progression were broadly repressed, consonant with our observation that both cell size and cell number are significantly lower in TRV::*NbPRS4* leaves. Genes involved in solute transport, nutrient uptake, and stress responses, including genes that encode heat shock proteins and glutathione S-transferases, are among the functional categories that were significantly induced in TRV::*NbPRS4*. The expression of various transcription factor (TF) families were also affected in TRV::*NbPRS4*, with a few strong patterns: TFs typically involved in developmental patterning and differentiation, such as the C2C2 Zn finger, MADS box, GRF, LOB, and homeobox TFs, were generally down-regulated in TRV::*NbPRS4*, whereas TFs typically involved in stress responses, such as ERF, NAC, bZIP, and WRKY TFs, were generally up-regulated in TRV::*NbPRS4* (Figure 4C). Thus, from a global perspective, the TRV::*NbPRS4* transcriptome demonstrates that loss of cytosolic PRPP severely disrupts cellular homeostasis, reprogramming the transcriptome to repress anabolic growth and developmental patterning and to induce a host of stress responses.

### Purine and pyrimidine biosynthesis drive TOR activity

PRS4-synthesized cytosolic PRPP contributes primarily to two pathways in plants cells: cytosolic PRPP is condensed with orotate by cytosolic UMP synthase (UMPSase), yielding orotidine 5′-monophosphate (OMP), a precursor in *de novo* pyrimidine biosynthesis; and adenine or hypoxanthine/guanine phosphoribosyltransferases convert adenine and guanine to AMP and GMP, respectively, using cytosolic PRPP in the purine salvage pathway. Since the TRV::*NbPRS4* knockdown drastically reduced TOR activity, we hypothesized that one or both of these pathways—*de novo* pyrimidine synthesis and/or purine salvage—are required to maintain TOR activity in plant cells.

To directly test this hypothesis, we expanded our genetic screen to silence genes that encode nucleotide biosynthesis enzymes in *N. benthamiana*. Specifically, we designed VIGS triggers to target critical genes in the *de novo* purine biosynthesis pathway (*PHOSPHORIBOSYLGLYCINAMIDE FORMYLTRANSFERASE*, or *GART*; *ADENYLOSUCCINATE LYASE*, or *ASL*; and *PHOSPHORIBOSYLAMINOIMIDAZOLE CARBOXAMIDE FORMYLTRASFERASE / INOSINE MONOSPHOSPHATE CYCLOHYDROLASE*, or *ATIC*) or in the *de novo* pyrimidine biosynthesis pathway (*DIHYDROOROTASE*, or *DHOase*; *DIHYDROOROTATE DEHYDROGENASE*, or *DHODH*; and *URIDINE MONOPHOSPHATE SYNTHASE*, or *UMPSase*). We selected these enzymes because they do not directly consume ATP or amino acids, and thus are specifically disrupting only nucleotide biosynthesis (and not incidentally impacting other primary metabolic pathways). Across experiments, silencing expression of enzymes in *de novo* pyrimidine or purine biosynthesis disrupted plant development and physiology, causing delayed growth, reduced leaf size, and striking chlorosis, among other phenotypes (Figure 5A). Strikingly, TOR activity was severely reduced after silencing genes involved in either *de novo* purine or pyrimidine biosynthesis, confirming our hypothesis that nucleotide biosynthesis is required to maintain TOR activity in plants (Figure 5B).

**Figure 5.**
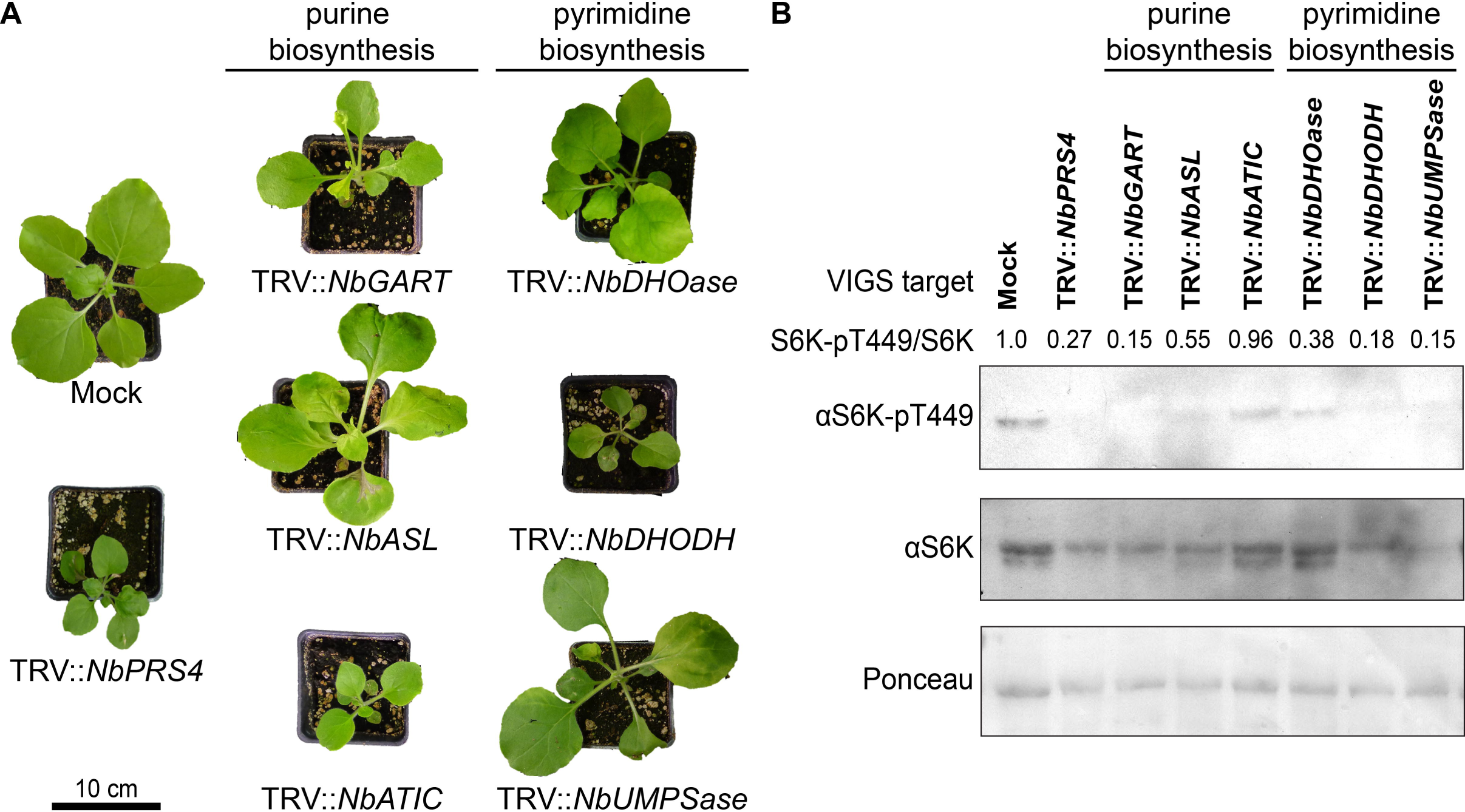
Silencing key genes in nucleotide biosynthesis inhibits TOR activity. (A) Nucleotide biosynthesis is necessary for normal shoot development and physiology. Silencing genes downstream of *PRS4* in nucleotide biosynthesis in *N. benthamiana* reduced leaf number and size, disrupted leaf shape, and caused chlorosis, similar to the phenotypes observed in TRV::*NbPRS4* plants. Each gene was silenced in at least six plants per experiment, and the entire experiment was replicated three times; representative individuals of each silenced gene are shown. (B) Silencing nucleotide biosynthesis genes lowers TOR activity. S6K-pT449 levels are strongly reduced in silenced plants compared to mock-infected controls, and the S6K-pT449/S6K ratios are consistently lower.

### Chemically inhibiting nucleotide biosynthesis inactivates TOR

We next tested whether the hypothesis that nucleotide biosynthesis drives TOR activity is conserved across plant species by investigating the effects of inhibiting nucleotide biosynthesis in *A. thaliana*. For this chemical genetic approach, we treated *A. thaliana* seedlings with inhibitors that specifically impact one or both of the PRS4-dependent nucleotide biosynthesis pathways, and then assayed for changes in TOR activity indicated by altered phosphorylation of S6K-T449. We screened four compounds that target processes that require cytosolic PRPP (Figure 6A). 5-fluorouracil (5FU) and 5-fluoroorotic acid (5FOA) limit only *de novo* pyrimidine synthesis by inhibiting UMPSase and thymidylate synthetase activity, respectively (Figure 6A). 6-mercaptopurine (6MP) specifically limits the purine salvage pathway by inhibiting PRPP transfer to purines by hypoxanthine phosphoribosyltransferases (Figure 6A). Methotrexate (MTX) limits both purine and pyrimidine biosynthesis by inhibiting dihydrofolate reductase, which is required for folate synthesis, a necessary nucleotide precursor (Figure 6A). To assay the effects of these inhibitors, seedlings were grown in ½ MS media for three days, then transferred to fresh ½ MS media supplemented with 15 mM glucose and 10 μM of one of the nucleotide biosynthesis inhibitors, or a mock treatment as a negative control. Seedlings were collected 24 h after treatment and proteins were extracted and analyzed by Western blot (Figure 6C). Under these conditions, MTX had the strongest effects. *De novo* pyrimidine synthesis inhibitors 5FU and 5FOA had less potent effects. 6MP, an inhibitor of the purine salvage pathway, had no noticeable effect on TOR activity, as reflected by S6K-pT449 levels. Some seedlings were left in media for 7 days after treatment to observe the inhibitors’ long-term effects on growth (Figure 6B). As in the western blot, MTX had the most severe effect on shoot and root growth (Figure 6B).

**Figure 6.**
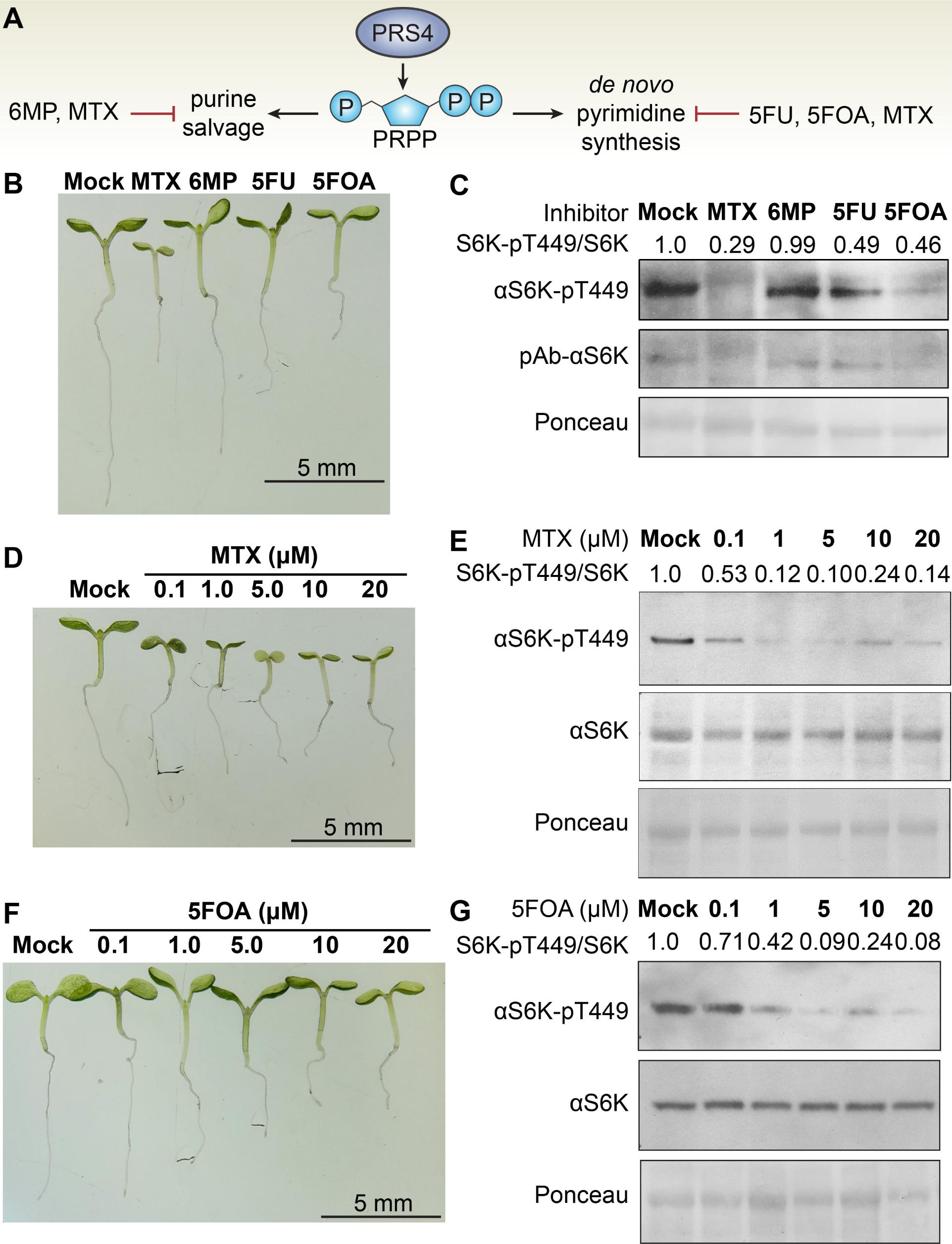
Chemically inhibiting nucleotide biosynthesis suppresses TOR activity. **(A)** PRPP synthesized by cytosolic PRS4 is primarily used for purine salvage and *de novo* pyrimidine synthesis. 6MP specifically inhibits purine salvage, 5FOA and 5FU specifically inhibit *de novo* pyrimidine synthesis, and MTX broadly inhibits nucleotide biosynthesis. **(B)** Seedlings were grown to quiescence, and then treated with either 15 mM glucose, which activates TOR and promotes root elongation (Xiong et al., 2013a), or 15 mM glucose and 10 μM MTX, 6MP, 5FU, or 5FOA, which limit nucleotide biosynthesis. Seedling growth was most drastically impaired in the presence of MTX, which inhibits all nucleotide biosynthesis. Treatment with 5FOA strongly impacted seedling growth as well, while 5FU and 6MP had less dramatic effects **(C)** Seedlings were treated with 10 μM of each inhibitor shown in panel A for 24 h. TOR activity was assayed using Western blots against S6K-pT449 and total S6K, with Ponceau staining shown as a loading control. MTX, 5FU, and 5FOA all reduced TOR activity. 6MP had no reproducible effect. This experiment was repeated three times; representative results are shown here. **(D)** Seedlings were grown as described in panel 6B and treated with either 0.1 μM, 1 μM, 5 μM, 10 μM, or 20 μM MTX. A photograph was taken 7 days after treatment. Growth was inhibited by all concentrations of MTX tested. **(E)** TOR activity was assayed using Western blots against S6K-pT449 and total S6K, with Ponceau staining shown as a loading control. 24-hour MTX treatment is effective at inhibiting TOR activity at a concentration as low as 0.1 μM, and1.0 μM MTX treatment effectively inactivates TOR entirely. **(F)** Seedlings were grown as described in panel 6B and treated with either 0.1 μM, 1 μM, 5 μM, 10 μM, or 20 μM 5FOA. A photograph was taken 7 days after treatment. 1 μM 5FOA was sufficient to noticeably delay seedling growth, and higher concentrations had more strong effects on inhibiting development. **(G)** TOR activity was assayed using Western blots against S6K-pT449 and total S6K, with Ponceau staining shown as a loading control. A 24-hour treatment with 5FOA has some effect at 0.1 μM, and strongly lowers TOR activity at concentrations ≥ 1.0 μM.

To demonstrate specificity and further investigate the effects of the nucleotide biosynthesis inhibitors that were most effective at reducing TOR activity (Figure 6C), we performed this assay again with a range of concentrations (0.1 μM, 1.0 μM, 5.0 μM, 10 μM, or 20 μM) of MTX and 5FOA. Consistent with our previous result, we found that MTX had the strongest effect, partially inhibiting growth and lowering S6K-pT449 levels even at a very low concentration of 0.1 μM. Treatment with 5.0 μM or more MTX caused nearly complete growth arrest, chlorosis, and an even greater decrease in S6K-pTT49 levels (Figure 6D and 6E). Low concentrations of 5FOA slightly reduced growth and had some impact on S6K-pT449 levels, whereas higher concentrations (5.0 μM to 20 μM) strongly inhibited growth and reduced S6K-pT449 levels (Figure 6F and 6G). Thus, we concluded that complete inhibition of *de novo* nucleotide synthesis by MTX was most effective at inhibiting TOR, inhibition of *de novo* pyrimidine synthesis was sufficient to lower TOR activity, and the purine salvage pathway was not needed to maintain TOR activity under these growth conditions. Given the potent impact of MTX on TOR activity, we focused further experiments on MTX treatments.

### Resupplying nucleotides restores TOR activity

First, we conducted a time course to determine how quickly MTX treatment impacts TOR activity. For these experiments, we used 0.1 μM MTX, which was an approximately minimal concentration to inhibit TOR activity 24 h after treatment. 2 h after MTX treatment, S6K-pT449 levels were not yet lower (Figure 7A). S6K-pT449 levels rapidly decreased over the next four hours, while total S6K levels remained stable (Figure 7A). Therefore, under our experimental conditions, MTX begins to lower TOR activity 3 to 4 h after treatment, ultimately abolishing S6K-pT449 levels by 24 h after treatment. This is consonant with past experiments using comparable experimental conditions, which showed that MTX requires at least ∼2-6 hours to begin to affect plant metabolism (Loizeau et al., 2008). To test the hypothesis that MTX inhibition of TOR activity is due to nucleotide depletion, we supplemented MTX-treated seedlings with 0.5 mM, 1.0 mM or 2.0 mM nucleotides: inosine monophosphates (IMPs), uracil monophosphates (UMPs), or a mixture of all nucleotide monophosphates (NMPs). We chose these concentrations because 1.0 – 2.0 mM nucleotides are a physiologically-relevant concentration (Chen et al., 2000). Neither IMPs nor UMPs were able to restore TOR activity after 3.5 h, in agreement with our results above suggesting that both purine and pyrimidine synthesis are required to maintain TOR activity in plants (Figure 7B and C). A mixture of all NMPs, however, was sufficient to significantly increase S6K-pT449 levels 3.5 h after treatment (Figure 7D). In summary, these experiments demonstrate that nucleotides are sufficient to activate TOR in MTX-treated seedlings. Therefore, we concluded that MTX inhibits TOR specifically by limiting nucleotide biosynthesis. Moreover, in agreement with our hypothesis that TRV::*NbPRS4* knockdowns reduce TOR activity due to limited nucleotide biosynthesis, these results directly demonstrate that TOR is sensitive to nucleotide availability in plants.

**Figure 7.**
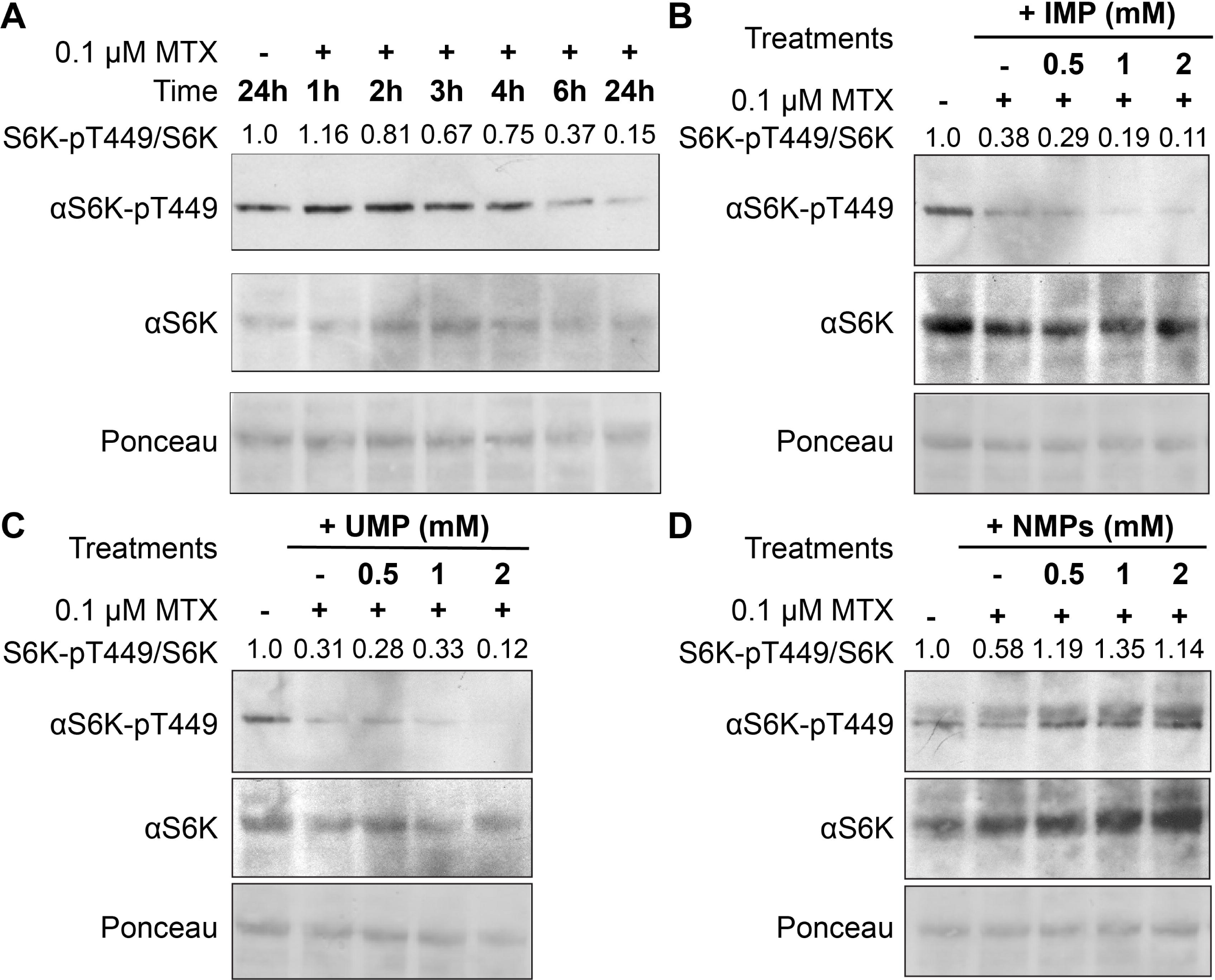
Nucleotide supply rescues TOR activity in MTX-treated seedlings. **(A)** Seedlings were treated with 0.1 μM MTX and collected over a time course, as indicated. TOR activity decrease 3 h after MTX treatment. This experiment was repeated three times; representative results are shown here. **(B,C,D)** Seedlings were pre-treated with 0.1 μM MTX for 3.5 h, and then supplied either inosine 5’-monophosphate (IMP) **(B)**, uracil 5’-monophosphate (UMP) **(C)**, or a mixture of nucleotide monophosphates (NMPs) **(D)** at physiologically-relevant concentrations (0.5, 1.0, or 2.0 mM). NMPs restored S6K-pT449 levels and S6K-pT449/S6K ratios in MTX-treated seedlings, suggesting that the effects of MTX on TOR activity are due to nucleotide depletion. Neither IMP nor UMP was sufficient to restore TOR activity, indicating that all nucleotides (purines and pyrimidines) are required to stimulate TOR in MTX-treated seedlings. This experiment was repeated five times; representative results are shown here.

### TOR promotes expression of *de novo* nucleotide biosynthesis genes

Among its many metabolic roles, TOR is known to promote nucleotide biosynthesis in animals and fungi through transcriptional and post-translational control (Ben-Sahra et al., 2013; Robitaille et al., 2013; Ben-Sahra et al., 2016). We therefore hypothesized that TOR could also transcriptionally promote nucleotide biosynthesis in plants. To test this hypothesis, we mined publicly available transcriptomic datasets, focusing on changes in expression of genes that encode nucleotide metabolic enzymes. We found that genes involved in nucleotide biosynthesis are enriched among the genes upregulated by the glucose-TOR transcriptional program (*p* < 0.01), as previously noted (Xiong et al., 2013a), and among the genes repressed when plants are treated with the TOR inhibitor AZD-8055 (*p* < 0.05) (Dong et al., 2015). Indeed, transcripts encoding enzymes of nearly every step of purine biosynthesis downstream from PRS-mediated PRPP synthesis are induced when TOR is activated and repressed when TOR is inactivated (Figure 8A). Several pyrimidine biosynthesis enzyme genes are also transcriptionally induced by TOR activation and transcriptionally repressed by TOR inactivation (Figure 8A). Moreover, genes encoding purine and pyrimidine salvage pathway enzymes and genes that promote DNA synthesis are also transcriptionally promoted by TOR (Supplemental Data Set 6). Alongside promoting nucleotide biosynthesis, TOR transcriptionally controls nucleotide catabolism: genes involved in P_i_ salvage, nucleotide degradation, and purine importers are all induced when TOR is inactivated, whereas genes involved in P_i_ salvage are repressed when TOR is activated. For instance, several NUDIX hydrolases (*NUDT8*, *NUDT13*, and *NUDT15*) are repressed ∼2-3.5-fold when TOR is activated by glucose (Supplemental Data Set 6). In concert, these data demonstrate that TOR activation broadly reprograms the plant transcriptome to promote nucleotide biosynthesis and repress nucleotide catabolism.

**Figure 8.**
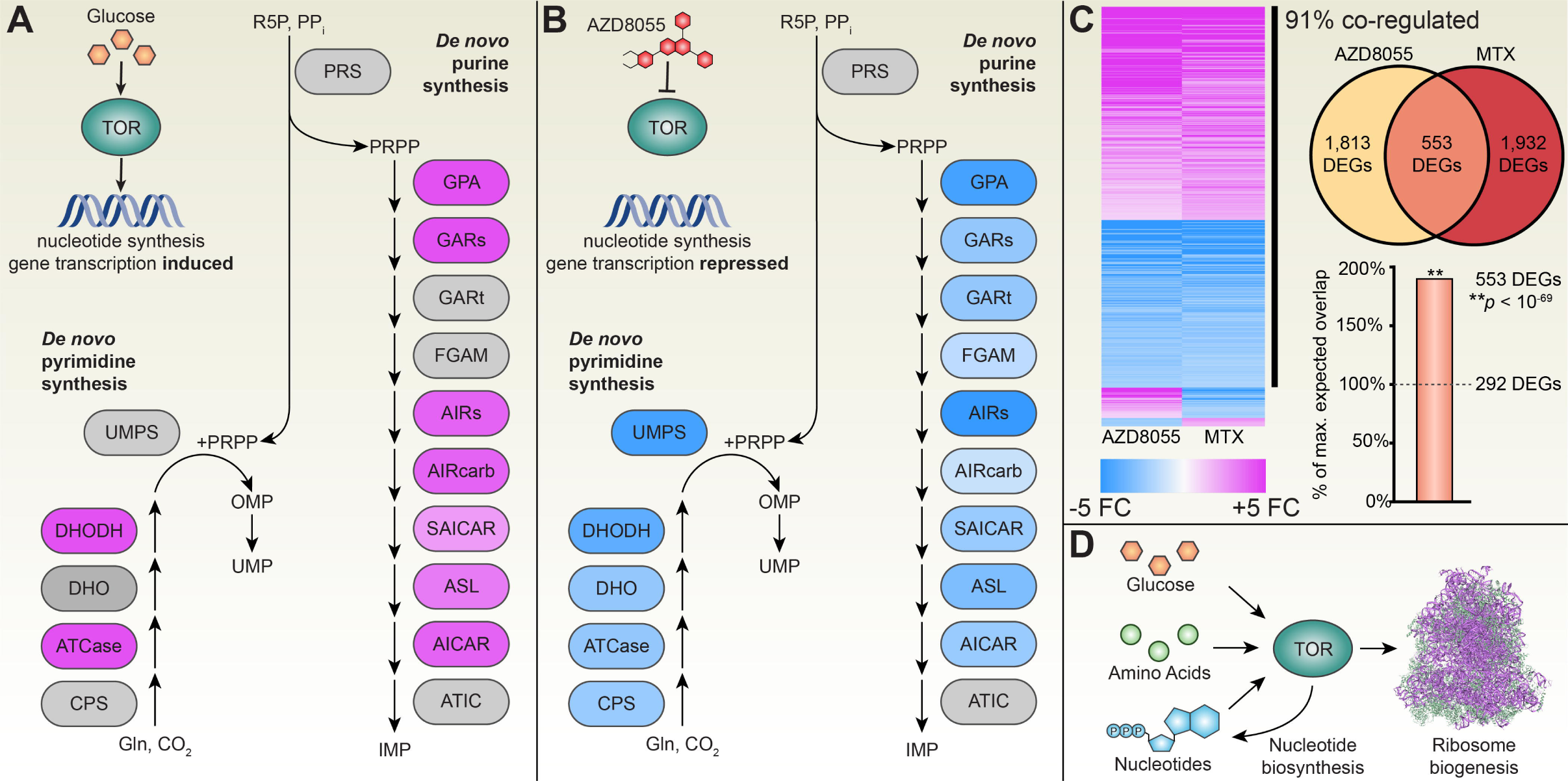
TOR coordinates nucleotide metabolism with ribosome biogenesis in plants. **(A,B)** Enzymes for each step of *de novo* pyrimidine (left) and *de novo* purine (right) biosynthetic pathways are shown. Fold-change in mRNA levels of these enzymes in (**A**) the glucose-TOR transcriptome (Xiong et al., 2013a) or (**B**) TOR-inhibited AZD8055 transcriptome (Dong et al., 2015) are indicated by color, as in panel C (induced genes are shown in magenta, repressed genes are shown in blue, grey indicates no detected difference in mRNA levels). **(C)** The set of significant DEGs after 24 h treatment with TOR inhibitor AZD8055 or nucleotide biosynthesis inhibitor MTX significantly overlap (*p* < 10^-69^), and 91% of overlapping DEGs are co-regulated. For details, see Supplemental Data Set 6. **(D)** Metabolites activate TOR to drive ribosome biogenesis. TOR integrates multiple metabolic signaling pathways in plants by sensing glucose, nucleotide, and (speculatively) amino acid levels. When other nutrients are available (e.g., upon glucose activation of TOR), TOR also drives nucleotide biosynthesis, which both reinforces TOR activity and metabolically supports rRNA synthesis to contribute to ribosome biogenesis.

We next reanalyzed the transcriptome of MTX-treated *A. thaliana* cells (Loizeau et al., 2008) to test how nucleotide deprivation impacts the *A. thaliana* transcriptional program. Using stringent cutoffs, our analysis revealed 16, 1,748, and 2,485 genes that are differentially expressed 2 h, 6 h, and 24 h after treatment with MTX, respectively, compared to mock-treated cells (Supplemental Data Set 6). The remarkably low number of DEGs detected 2 h after MTX treatment is in keeping with our conclusion above that MTX treatment has effectively no impact on TOR activity 2 h after treatment. Therefore, we focused our analysis on the other two timepoints, 6 h and 24 h after MTX treatment. The biological functions of the DEGs 6 h and 24 h after MTX treatment are largely similar. Broadly, MTX treatment represses expression of genes involved in ribosome biogenesis, mitochondrial oxidative phosphorylation (OXHPOS), and photosynthesis, while inducing expression of E3 ubiquitin ligases that promote protein degradation and amino acid recycling, WRKY and AP2/EREBP transcription factors involved in stress responses, and various starvation response genes, e.g., the *DARK-INDUCIBLE* (*DIN*) genes *DIN2*, *DIN6*, and *DIN10* (Supplemental Data Sets 6 and 7). As with the TRV::*NbPRS4* transcriptome, many of these gene categories are conserved targets of TOR, such as ribosomal protein genes and mitochondrial OXPHOS components, which are consistently induced by TOR activity throughout eukaryotic lineages. Indeed, we found that the set of genes regulated by 24 h treatment with AZD-8055, a highly specific ATP-competitive TOR kinase inhibitor, significantly overlapped with the set of genes regulated by 24 h treatment with MTX (553 genes, *p* < 10^-69^; for comparison, the maximum expected overlapping genes is 292, *p* = 0.05), and that these overlapping genes are almost all co-regulated (503 genes, or 91%, are either induced or repressed in both MTX- and AZD-8055 transcriptomes) (Figure 8C, Supplemental Data Set 6). The striking similarity between the transcriptional profiles of MTX- and AZD-8055-treated *A. thaliana* demonstrates that many of the effects of MTX-mediated nucleotide depletion can be reproduced by TOR inhibition, indicating that nucleotide sensing by TOR is a crucial signaling axis to maintain metabolic homeostasis (Figure 8D).

## Discussion

### Functional genetics to probe TOR signaling networks

TOR is an essential regulatory kinase across all lineages of eukaryotes (Chantranupong et al., 2015). In *A. thaliana*, null *tor* alleles cause embryo arrest at early stages of embryogenesis (Menand et al., 2002). Chemically inhibiting TOR kinase activity or genetically silencing *TOR* expression causes plant growth arrest at later stages of development (Ahn et al., 2011; Montané and Menand, 2013; Xiong et al., 2013b). Therefore, we hypothesized that loss of genes required for activity would likely also be embryo-lethal. To circumvent conducting an arduous forward genetic screen of embryo-lethal mutants (Kim et al., 2002), we took a “fast reverse genetics” approach (Baulcombe, 1999), silencing predicted embryo-lethal genes in *N. benthamiana* during vegetative development with virus-induced gene silencing. With this approach, we discovered that plant TOR activity is sensitive to nucleotide availability in plants, consonant with the recent discovery that mammalian TOR also senses nucleotide availability (Hoxhaj et al., 2017; Emmanuel et al., 2017).

Silencing embryo-lethal genes with VIGS does not always cause severe developmental phenotypes (Stonebloom et al., 2009; Burch-Smith and Zambryski, 2010; Burch-Smith et al., 2011), but silencing the cytosolic phosphoribosyl pyrophosphate synthetase, *PRS4*, dramatically disrupted plant growth and development. It should be noted that TRV::*NbPRS4* knockdown phenotypes are more severe than TRV::*NbTOR* knockdown phenotypes (Ahn et al., 2011), suggesting that loss of cytosolic PRPP does not exclusively arrest growth via inhibition of the TOR signaling pathway. Similarly, homozygous *prs4-1* knockout embryos arrest at the very earliest stages of embryogenesis, whereas homozygous *tor* knockout embryos arrest at the globular stage. We speculate that the more severe phenotypes of TRV::*NbPRS4* knockdowns and knockouts reflect its absolute requirement for nucleotide biosynthesis to support DNA synthesis during cell division.

### TOR coordinates nucleotide metabolism with ribosome biogenesis

In plant leaves, 40-60% of P_o_ is incorporated in ribosomal RNA (rRNA). rRNA expression and stability are directly controlled by TOR, which promotes expression of ribosomal protein genes, rRNA, and other genes involved in ribosome biogenesis (Ren et al., 2011; Kim et al., 2014). In mutant human cell lines with high TOR activity, overproduction of rRNA disrupts nucleotide homeostasis, making nucleotides less available for other biological processes (Valvezan et al., 2017). As a consequence, these cell lines are hypersensitive to nucleotide synthesis inhibitors, which rapidly deplete nucleotide availability for DNA replication, ultimately triggering DNA damage responses and cell death (Valvezan et al., 2017). Here, we have shown that TOR similarly coordinates nucleotide supply and demand in plant cells. TOR promotes transcription of genes involved in nucleotide biosynthesis and ribosome biogenesis. If nucleotide levels decrease due to another nutritional limitation or physiological stress, i.e., despite the transcriptional promotion of nucleotide biosynthesis by TOR, TOR activity decreases, reducing the demand for nucleotides by ribosome biogenesis. Based on our findings, and by analogy to the human nucleotide-TOR signaling pathway (Valvezan and Manning, 2019), we propose that TOR acts as a molecular rheostat to maintain nucleotide homeostasis in plant cells (Figure 6D).

The mechanism of nucleotide sensing by TOR in plants remains unresolved by this study. In human cells, nucleotide depletion is currently proposed to act through the TSC-Rheb signaling axis to inhibit TOR activity (Emmanuel et al., 2017; Hoxhaj et al., 2017). Rheb is a small GTPase that activates TOR at the lysosomal surface, and Rheb engages TOR when growth factor signals suppress its GTPase-activating protein complex, TSC. In nucleotide-depleted cells, Rheb GTP-loading decreases, and its localization, stability, and activity are compromised (Emmanuel et al., 2017; Hoxhaj et al., 2017). Neither Rheb nor TSC are deeply conserved in eukaryotic lineages, so plants must have evolved a distinct mechanism for sensing nucleotide availability. Identifying the nucleotide sensor will be aided by ongoing efforts to elucidate the plant TOR signaling network (Wu et al., 2019).

### TOR and Po sensing for sustainable agriculture

The TOR signaling network is a promising candidate for improving agricultural crops. Current agricultural systems are unsustainable, but biotechnological enhancement of crops can make significant contributions to sustainability efforts (Springmann et al., 2018). To date, most studies of nutrient sensing in plants have focused on pathways that detect inorganic nutrients, such as nitrate, ammonium, and phosphate (Chiou and Lin, 2011; Xu et al., 2012). In natural settings, however, plants consume both inorganic and organic nutrients, such as dissolved P_o_ and amino acids (Näsholm et al., 1998). Moreover, once nutrients are absorbed by plants, many species rapidly convert inorganic nitrogen and phosphate to organic forms for transport and metabolism. Therefore, understanding the mechanisms of organic nutrient sensing in plants, including the TOR signaling network, should accelerate efforts to develop sustainable agricultural crop varieties.

Here, we showed that TOR is sensitive to nucleotide availability, a major fraction of the P_o_ available for anabolism (Figure 1A, 8D). Moreover, TOR activity strongly promotes synthesis of nucleotides and ribosomal nucleic acids to support the high ribosome abundance required for protein translation in metabolically-active cells. Accordingly, genetically or chemically dampening TOR activity could reduce phosphorus demands, by drastically decreasing the rate of nucleotide and nucleic acid synthesis. Alternatively, for plants grown with excess phosphorus supply, stimulating TOR activity could increase growth rates and crop yields by increasing phosphorus use efficiency and allocation of phosphorus to bioactive, productive macromolecules (nucleotides and rRNA). Therefore, rewiring the TOR signaling network in crop species could theoretically contribute to enhancing crop yields and reducing reliance on fertilizers.

## Methods

### Plant materials and growth conditions

Plants were grown under standard conditions with 16 h day / 8 h night at ∼120 µmol photons m^-2^ s^-1^ of photosynthetically active radiation and at 22°C-24°C unless otherwise stated. The inbred Col-0 ecotype was used as wild-type for all *A. thaliana* seedling experiments. GabiKat-780B11, which had been previously described (Bolle et al., 2013), was obtained from the Arabidopsis Biological Resources Center. The reference inbred Nb-1 ecotype, obtained from the Boyce Thompson Institute, was used for all *N. benthamiana* experiments. *prs4-1* was genotyped by extracting DNA from Col-0 plants by grinding snap-frozen leaf tissue in DNA extraction buffer (200 mM Tris-Cl at pH 5.7, 250 mM NaCl, 25 mM EDTA, 0.5% SDS), precipitating DNA in isopropanol, washing DNA with 70% ethanol, and resuspending purified DNA in ddH_2_O. PCRs were used to assay for wild-type or T-DNA insertion alleles in 20 μL reactions using GoTaq Green Master Mix (Promega) using manufacturer’s instructions and 30 cycles as follows: 95°C for 30 seconds, 57°C for 30 seconds, and 72°C for 60 seconds. Genotyping primers used were: 5′-CAA GGA TTG TCT CTA ATA TCC CCA-3′ (left genomic primer, LP), 5′-CTT TGG GAA GAC ACC ATG AGT TAC-3′ (right genomic primer, RP), and 5′-ATA ATA ACG CTG CGG ACA TCT ACA TTT T-3′ (T-DNA border primer, LB).

### Molecular cloning

Virus-induced gene silencing vectors were prepared as previously described (Brunkard et al., 2015). Briefly, RNA was isolated from Nb-1 shoots with the Spectrum Plant Total RNA kit (Sigma-Aldrich) with on-column DNase I digestion (New England Biolabs). cDNA was synthesized from RNA using random hexamers and SuperScript III reverse transcriptase (Fisher Scientific). The *NbPRS4* silencing trigger was amplified with Phusion DNA polymerase (New England Biolabs) using oligonucleotides 5′-GCA TCT AGA ATG GAG AAT GGT GCG C-3′ and 5′-GAT CTC GAG TTC AAC AGT GGG ATA CC-3′ as primers. The amplified *NbPRS4* PCR product was gel purified (Monarch DNA Gel Extraction Kit, New England Biolabs), digested with XbaI and XhoI (New England Biolabs), and ligated with XbaI- and XhoI-digested pYL156 (Liu et al., 2002) using T4 DNA ligase (Promega). Ligations were transformed into house-made chemically competent *E .coli* DH10B. The silencing triggers for all other nucleotide biosynthesis genes were amplified with Phusion DNA polymerase (New England Biolabs) using the oligonucleotides listed in Supplemental Data Set 8. The amplified products were gel purified (Monarch DNA Gel Extraction Kit, New England Biolabs) and incubated with 100 ng XbaI- and XhoI-digested pYL156 in a molar ratio of 1 vector : 2 insert. Reactions were incubated with T4 DNA polymerase (New England Biolabs) in NEBuffer 2.1 (New England Biolabs) for 2.5 minutes at room temperature, placed on ice for 10 minutes, and then transformed into house-made chemically competent *E .coli* DH10B. Plasmids were then miniprepped with Bioneer kits, Sanger sequenced to confirm insert sequences, and transformed into *Agrobacterium* GV3101. Manufacturer’s protocols were followed throughout.

To clone *AtPRS4* for complementation experiments, cDNA was synthesized from RNA isolated from Col-0 rosette leaves, using the techniques described above. *PRS4* was amplified by PCR using oligonucleotides 5′-cacc ATG TCT GAG AAC GCA GCC A-3′ and 5′-AAT CTG CAG AGC ATC AGC AAT-3′ and ligated with topoisomerase into a Gateway entry vector (pENTR/D-TOPO, ThermoFisher). This vector was then digested with EcoRV (New England Biolabs) and recombined with pEarleyGate expression vectors pEarleyGate103 and pEarleyGate104 (Earley et al., 2006) using LR Clonase II (ThermoFisher). After the sequence of these vectors was confirmed, each was transformed into *Agrobacterium* GV3101 for subsequent transformation of plants.

### TOR activity assays

*A. thaliana* seedlings or *N. benthamiana* leaves were snap-frozen in liquid nitrogen. Protein was then extracted from the plant tissue in 100 mM MOPS (pH 7.6), 100 mM NaCl, 5% SDS, 0.5% β-mercaptoethanol, 10% glycerin, 2 mM PMSF, and 1x PhosSTOP phosphatase inhibitor (Sigma-Aldrich). S6K-pT449 was detected by Western blot using a phosphospecific antibody (ab207399, AbCam) and an HRP-conjugated goat anti-rabbit IgG secondary antibody (Jackson Immuno Research, no. 111-035-003). S6K levels were detected by Western blot using a custom monoclonal antibody described below. Total protein was visualized after transfer using Ponceau S red staining. Western blot images were scanned, converted to grayscale, and adjusted for contrast and brightness using ImageJ.

To generate monoclonal antibodies that detect total S6K protein levels, peptides were synthesized that corresponded to amino acids 439 through 459 surrounding the TOR substrate T449 in AtS6K1 (At3g08730.1) with either threonine or phosphothreonine at T449 (DPKANPFTNFTYVRPPPSFLH or DPKANPFTNFpTYVRPPPSFLH) and conjugated to either BSA or KLH (GenScript). BSA-conjugated phosphopeptide was used to immunize mice. Sera were screened with enzyme-linked immunosorbet assays (ELISAs) for reactivity to KLH-conjugated peptide and/or phosphopeptide. Subsequent screening of hybridomas with ELISAs using KLH-conjugated peptide or phosphopeptide and Western blots against S6K protein identified a monoclonal antibody that reliably detects total S6K protein levels, regardless of phosphorylation status of T449. Throughout, we refer to this monoclonal antibody as αS6K. In one preliminary experiment (Fig. 6C), we used a commercially-available polyclonal antibody that detects total S6K levels (sc-230, Santa Cruz Biotechnology); we refer to this antibody as pAb-αS6K.

### Virus-induced gene silencing

VIGS was performed as previously described (Brunkard et al., 2015). Briefly, *Agrobacterium* cultures carrying either pYL156 containing the silencing trigger or pYL192 were grown overnight in LB with antibiotics at 28°C. Cultures were spun down at 700 × *g* for 10 minutes, washed, and resuspended to OD_600nm_ = 1.0 in 10 mM MgCl_2_, 10 mM 2-(*N*-morpholino)ethanesulfonic (MES) acid (pH = 5.7, adjusted with KOH), and 200 µM acetosyringone from a recently-prepared 100mM stock solution frozen in DMSO. Agrobacteria were left to induce virulence at room temperature for 2-4 h with gentle shaking before pressure infiltration into 3-week old *N. benthamiana* leaves by needleless syringe. Plants were returned to standard growth conditions and monitored. Tissue collection and phenotypic observation were performed 2 weeks post-infiltration. TRV::*NbPDS* was used as a positive control to track viral infection and VIGS efficiency, and TRV::*GUS* (pYC1) was used as a mock treatment negative control (Stonebloom et al., 2009). Each gene was silenced in at least 6 plants per experiment. All VIGS experiments were performed at least 3 times.

### Phenotypic analysis of TRV::*NbPRS4* plants

All phenotypic analyses and photography of TRV::*NbPRS4* plants were performed 2 weeks post-agroinfiltration. Shoot apices were collected for histology and SEM analysis. The 4^th^ youngest leaf (i.e., the 4^th^ leaf counting down from the apex of each plant) was used for quantifying epidermal cell size. Forceps were used to peel off sections of epidermis from the adaxial side of the leaf, which was then observed under a compound light microscope with a 40x objective. Cells were outlined manually in ImageJ to determine area in square micrometers. Measurements were performed on GUS (“mock”) and “moderate” TRV::*NbPRS4* individuals, as tissue from “severe” plants was too fragile to process. Plants unused in other experiments were allowed to grow indefinitely so leaf initiation rate, leaf morphology, and flowering time could be observed. To quantify rRNA levels, total RNA was extracted using the Sigma Spectrum™ Plant Total RNA kit and analyzed using the Agilent RNA 6000 Pico Chip with the Agilent Bioanalyzer 2100.

For histology, meristems were harvested from *N. benthamiana* two weeks after inoculation with *Agrobacterium* cultures carrying either pYL156::NbPRS4 or pYL156::GUS (pYC1). Meristems were vacuum infiltrated with infiltration solution (50% ethanol, 5.0% acetic acid, 3.7% formaldehyde) at 15 Hg for 15 minutes. Tissue was dehydrated through a stepwise ethanol dehydration series (15%, 20%, 50%, 75%, 95%) with 15-minute incubations at each step and stored overnight in 95% ethanol + 0.1% Eosin solution. Samples were washed for 20 minutes with the following solutions: 100% ethanol (twice), 75% ethanol + 25% Histoclear, 50% ethanol + 50% Histoclear, 25% ethanol + 75% Histoclear, 100% Histoclear (twice). Tissue was stored overnight in 1:1 Histoclear : melted paraplast solution. The next morning, Histoclear : paraplast solution was removed and replaced with melted paraplast. Samples were incubated at 56°C for 8-10 hours. Paraplast was replaced every 8-10 hours six additional times. Samples were poured into plastic weighboats, allowed to harden, sectioned, and mounted onto slides. Slides were rehydrated through an ethanol series (100%, 100%, 95%, 85%, 70%, 50%, 30%, 15%), stained in 0.1% toluidine blue, and then partially dehydrated through an ethanol series (15%, 30%, 50%, 70%) to destain slightly. Slides were hydrated again (70%, 50%, 30%, 15%) and rinsed in distilled water. Mounted tissue was treated with ImmunoHistoMount and covered with a cover slip. Meristems were visualized with a compound microscope.

### Phenotypic analysis of *prs4-1* mutants

*prs4-1 /* + heterozygous *A. thaliana* mutants were allowed to self-fertilize for analysis of homozygous *prs4-1 / prs4-1* progeny in siliques. The location of homozygous *prs4-1* / *prs4-1* progeny along siliques was marked following standard protocols (seedgenes.org). For positional analysis, the number of aborted seeds in the first ten seeds from the silique base (i.e., seeds #1 through #10) and in the second ten seeds from the silique base (i.e., seeds #11 through #20) was counted in nine independent siliques at comparable developmental stages from several independent *prs4-1* / + parents. Seed abortion was not observed in these positions in wild-type siblings grown under the same conditions. Chi-square tests were used to compare the frequency of aborted seeds in these positions to the expected frequency (25%) of aborted seeds. To visualize embryos in *prs4-1* / + progeny, *A. thaliana* seeds were removed from young siliques and placed on one drop of Hoyer’s solution on a microscope side. Embryos were visualized with a compound microscope after one hour of clearing in Hoyer’s solution.

### RNA-Seq and transcriptome analysis

Tissue for RNA-Seq was collected from TRV::*NbPRS4* or mock-treated plants 2 weeks post-agroinfiltration with TRV binary vectors. Plant apices, including the youngest 2-3 leaves, were collected and immediately frozen in liquid nitrogen. 9 mock-treated individuals, 9 TRV::*NbPRS4* individuals that showed “severe” phenotypic abnormalities, and 9 “moderate” TRV::*NbPRS4* individuals were collected. RNA was extracted using Sigma Spectrum™ Plant Total RNA kit. 3 pools of RNA from 3 individuals were prepared for the mock, severe, and moderate collections. RNA pools were made by combining 333.3 ng of RNA from each individual. Illumina TruSeq Stranded Total RNA kit with Ribo-Zero Plant was used to prepare libraries for RNA sequencing. Libraries were sequenced at the Vincent J. Coates Genomics Laboratory (QB3) using the Illumina Hi-Seq 4000. Reads were aligned with HISAT2 (Kim et al., 2015) and counted with HTSeq (Anders et al., 2015). Differential transcript abundance was determined with DESeq2 (Love et al., 2014).

The methotrexate-treated transcriptome was previously described (Loizeau et al., 2008); briefly, cells were treated with 100 μM methotrexate or mock-treated for 2 h, 6 h, or 24 h, RNA was extracted, and cRNA was hybridized with the CATMA array to detect changes in transcript levels. Only transcripts that were detected at significantly different levels in both treatment replicates (*p* < 0.05) were considered significant DEGs here.

*PRS* expression profiles during plant development were obtained from the Bio-Analytic Resource for Plant Biology (bar.utoronto.ca) with expression data and an electronic Fluorescent Pictograph (eFP) developed for AtGenExpress (Schmid et al., 2005).

Transcriptomes were further analyzed with MapMan software (Thimm et al., 2004). The *N. benthamiana* genome was annotated for gene function using Mercator4 (Lohse et al., 2014). Significantly affected gene categories were determined by MapMan using a Wilcoxon rank-sum test with the Benjamini-Hochberg-Yekutieli procedure to correct for the false discovery rate. All MapMan annotations and statistical analyses are available for review in supplemental data. The lists of differentially expressed genes from AZD-8055-treated and MTX-treated samples were compared using a hypergeometric test for significant overlap using R software (phyper function). For comparative analysis, only those genes that could be detected in both the MTX and AZD-8055 treatment transcriptomes (i.e., the ∼22,000 genes with complementary probes in the CATMA microarray) were considered.

### Seedling chemical treatment assays

*35S_PRO_:S6K1-HA* Col-0 seeds (Xiong and Sheen, 2012) were sterilized with a 50% bleach and 0.1% tween-20 solution and stratified in the dark at 4°C for 2 days in ddH_2_O. Each well of a standard 6-well culture plate was filled with 1 mL of ½ MS, pH 5.7liquid media. ∼20 *35S_PRO_:S6K1-HA* Col-0 seeds were placed in each well under aseptic conditions, and the plate was sealed with micropore tape. Plates were grown at 23°C with a light intensity of 70 μmol m^−2^ s^−1^ photosynthetically active radiation and a 12 h day / 12 h night cycle for 3 days. The media were then replaced with ½ MS supplemented with 15 mM D-glucose, pH 5.7, and any other treatments as described in the text, and returned to growth chambers. All treatments, transfers, and seedling collections were performed at subjective dawn unless otherwise noted for consistency. Photographed seedlings were left in treatment media for 7 days to observe developmental differences.

Methotrexate (Gainger), 6-mercaptopurine (Fisher Scientific), 5-fluorouracil (Fisher Scientific), 5-fluoroorotic acid (Fisher Scientific), adenosine-5′-monophosphate disodium (Fisher Scientific), cytidine-5′-monophosphate disodium (Chem-Impex), guanosine-5′-monophosphate disodium (Chem-Impex), inosine-5’-monophosphate (TGI), and uracil 5′-monophosphate trisodium (Chem-Impex) were prepared fresh in concentrated stock solutions that were then diluted in ½ MS media (Caisson Labs), pH = 5.7 with KOH, for application to seedlings.

## Acknowledgments

This project was supported by NIH DP5-OD-023072 to J.O.B. M.B. is supported by an NSF Graduate Research Fellowship. This work used the Vincent J. Coates Genomics Sequencing Laboratory at UC Berkeley, supported by the NIH-S10-OD018174 Instrumentation Grant. We thank De Wood and Tina Williams at USDA ARS for microscopy support, Prof. Jen Sheen for sharing *35S_PRO_:S6K-HA A. thaliana* seeds, Anjuli Matharu for experimental assistance, and Sam Leiboff for assistance with RNA-Seq analysis.

## Author Contributions

M.B., M.R.S., R.H., and J.O.B. designed the research, performed experiments, and analyzed data. M.B. and J.O.B. wrote the manuscript.

---

**Supplemental Figure 1. P*R*S4 is expressed throughout plant development.** *PRS4* expression shown in the *A. thaliana* eFP browser (top panel); magenta indicates high relative transcript abundance, white indicates low relative transcript abundance. *PRS4* is most strongly expressed in metabolically active developmental stages, such as imbibed (germinating) seeds and shoot apices. Expression patterns of all *PRS* genes, except for *PRS1*, are shown in a bar graph (bottom panel). *PRS4* is consistently expressed at higher levels that the other *PRS* genes, except for during late seed development and in mature pollen.

**Supplemental Figure 2. (A)** RNA was extracted from shoot apices of *N. benthamiana* plants as shown. **(B)** Schematic of seedling treatment assays. Briefly, after stratification, seeds were germinated at subjective dawn in ½ MS and allowed to grow for three days (to quiescence). Seedling media was then replaced with ½ MS and 15 mM D-glucose, and left for 24 h to activate TOR and initiate growth. Then, seedlings were treated with 0.1 μM MTX or other nucleotide biosynthesis inhibitors, and collected as described in the text. Alternatively, after 24 h treatment with MTX, plants were re-supplied with nucleotide monophosphates,, or mock treatments, and collected for analysis.

**Supplemental Data Set 1.** Significant DEGs in TRV::*NbPRS4* knockdown *N. benthamiana*.

**Supplemental Data Set 2.** Complete transcriptome of TRV::*NbPRS4* knockdown *N. benthamiana*.

**Supplemental Data Set 3.** Significant DEGs in moderate versus severe TRV::*NbPRS4* knockdown *N. benthamiana*.

**Supplemental Data Set 4.** Mercator4 annotation of the Nb-1 v1.0.1 transcriptome.

**Supplemental Data Set 5.** MapMan functional analysis of the TRV::*NbPRS4* transcriptome.

**Supplemental Data Set 6.** Transcriptomes of *A. thaliana* treated with MTX, AZD8055, or glucose-TOR treatments.

**Supplemental Data Set 7.** MapMan functional analysis of the MTX transcriptome.

**Supplemental Data Set 8.** Oligonucleotides used in this study.

## Notes

#### Summary of Updates

Author list updated.

